# miR-17∼92 exerts stage-specific effects in adult V-SVZ neural stem cell lineages

**DOI:** 10.1101/2021.07.28.454109

**Authors:** Fabrizio Favaloro, Annina DeLeo, Ana C. Delgado, Fiona Doetsch

**Affiliations:** Biozentrum, University of Basel, CH 4056 Basel, Switzerland; Department of Pathology and Cell Biology, Columbia University, New York, NY 10032, USA

**Keywords:** miR-17∼92, V-SVZ, neural stem cells, adult neurogenesis, oligodendrogenesis, gliogenesis, intraventricular OPCs

## Abstract

In the adult mouse brain, neural stem cells (NSCs) in the ventricular-subventricular zone (V-SVZ) generate neurons and glia throughout life. microRNAs are important regulators of cell states, frequently acting in a stage- or context-dependent manner. Here, miRNA profiling of FACS-purified populations identified miR-17∼92 as highly upregulated in activated NSCs and transit amplifying cells (TACs) in comparison to quiescent NSCs. Conditional deletion of miR-17∼92 in NSCs reduced stem cell proliferation both *in vitro* and *in vivo*. In contrast, in TACs, miR-17∼92 deletion caused a selective shift from neurogenic DLX2^+^ TACs towards oligodendrogenic OLIG2^+^ TACs, resulting in increased oligodendrogenesis to the *corpus callosum*. miR-17∼92 deletion also decreased proliferation and maturation of intraventricular oligodendrocyte progenitor cells. Together, these findings reveal stage- and cell-type-specific functions of the miR-17∼92 cluster within adult V-SVZ neural stem cell lineages.

## Introduction

In the adult mouse brain, the V-SVZ, adjacent to the lateral ventricles, is the largest germinal niche. V-SVZ NSCs are radial cells that express glial fibrillary acidic protein (GFAP). They are largely quiescent (qNSCs) and, upon activation (aNSCs), give rise to transit amplifying cells (TACs), which in turn generate neuroblasts that migrate to the olfactory bulb (OB) (Chaker et al., 2016; Obernier and Alvarez-Buylla, 2019). Importantly, NSCs also give rise to a small number of glial cells, including oligodendrocytes destined for the *corpus callosum* and intraventricular oligodendrocyte progenitor cells (OPCs) (Chaker *et al*., 2016; Delgado et al., 2021; Obernier and Alvarez-Buylla, 2019).

Cell transitions along the stem cell lineage are underlain by key changes in gene regulatory networks. Among the players orchestrating progression along the lineage, microRNAs (miRNAs) are emerging as important regulators for their ability to rapidly sculpt cell transcriptomes. miRNAs are a class of small non-coding RNAs that are able to modulate cell states by targeting hundreds of mRNA transcripts simultaneously, either repressing their translation or promoting their degradation (O’Brien et al., 2018). To date, the regulation of adult NSC behavior by miRNAs is still little explored. Indeed, only a handful of miRNAs have been implicated in OB neurogenesis so far (Akerblom et al., 2014; Brett et al., 2011; Cheng et al., 2009; de Chevigny et al., 2012; Lepko et al., 2019; Liu et al., 2010; Pathania et al., 2012; Szulwach et al., 2010; Zhao et al., 2010; Zhao et al., 2009). Therefore, to expand the repertoire of miRNAs relevant to early lineage transitions in the V-SVZ, we performed miRNA profiling of FACS-purified qNSCs, aNSCs and TACs and found the miR-17∼92 cluster to be significantly upregulated in aNSCs and TACs in comparison to qNSCs.

miR-17∼92 is a cluster of six co-transcribed miRNAs (−17, -18a, -19a and b, -20a, and -92a) with pleiotropic functions in several tissues during both development and tumorigenesis (Concepcion et al., 2012). During embryogenesis, miR-17∼92 promotes the expansion of radial glial cells concomitantly preventing their premature transition into intermediate progenitors (Bian et al., 2013), and sustains the neurogenic phase by inhibiting acquisition of gliogenic competence (Naka-Kaneda et al., 2014). In the adult V-SVZ, miR-17∼92 is up-regulated upon stroke where it induces proliferation and survival of progenitor cells (Liu et al., 2013). However, its role under homeostasis is unknown.

Here, we show that miR-17∼92 is expressed in the adult V-SVZ neurogenic lineage and that its deletion results in stage-specific effects. In NSCs, miR-17∼92 deletion reduces proliferation while in TACs, it causes a shift from neurogenic to oligodendrogenic progenitors, thereby increasing oligodendrogenesis to the *corpus callosum*. Finally, in intraventricular OPCs, miR-17∼92 deletion impairs proliferation and maturation. Altogether, our results unveil new functions of the miR-17∼92 cluster in stage-specific regulation within V-SVZ lineages.

## Results

### miR-17∼92 is expressed in the adult V-SVZ neurogenic lineage

qNSCs, aNSCs and TACs can be directly purified from the adult V-SVZ niche using fluorescence activated cell sorting (FACS) (Codega et al., 2014; Pastrana et al., 2009) (Suppl. Fig. 1A). To identify miRNAs relevant to the early stages of the V-SVZ stem cell lineage we performed miRNA profiling of FACS-purified populations using Taqman miRNA arrays (Table S1). This analysis revealed cohorts of miRNAs enriched in qNSCs, aNSCs and TACs. Interestingly, all miRNAs significantly enriched in aNSCs over qNSCs were members of the miR-17∼92 cluster and its paralogs, miR-106a∼363 and miR106b∼25 (Suppl. Fig. 1B), and were also expressed in TACs (Fig 1A). While miR-106b∼25 promotes adult V-SVZ neural stem/progenitor cell proliferation and differentiation *in vitro* (Brett *et al*., 2011), it is unknown whether the miR-17∼92 and miR-106a∼363 clusters regulate the adult V-SVZ under homeostasis. Here, we focused on the role of the miR-17∼92 cluster in the regulation of early stages of the adult V-SVZ lineage.

**Figure 1.**
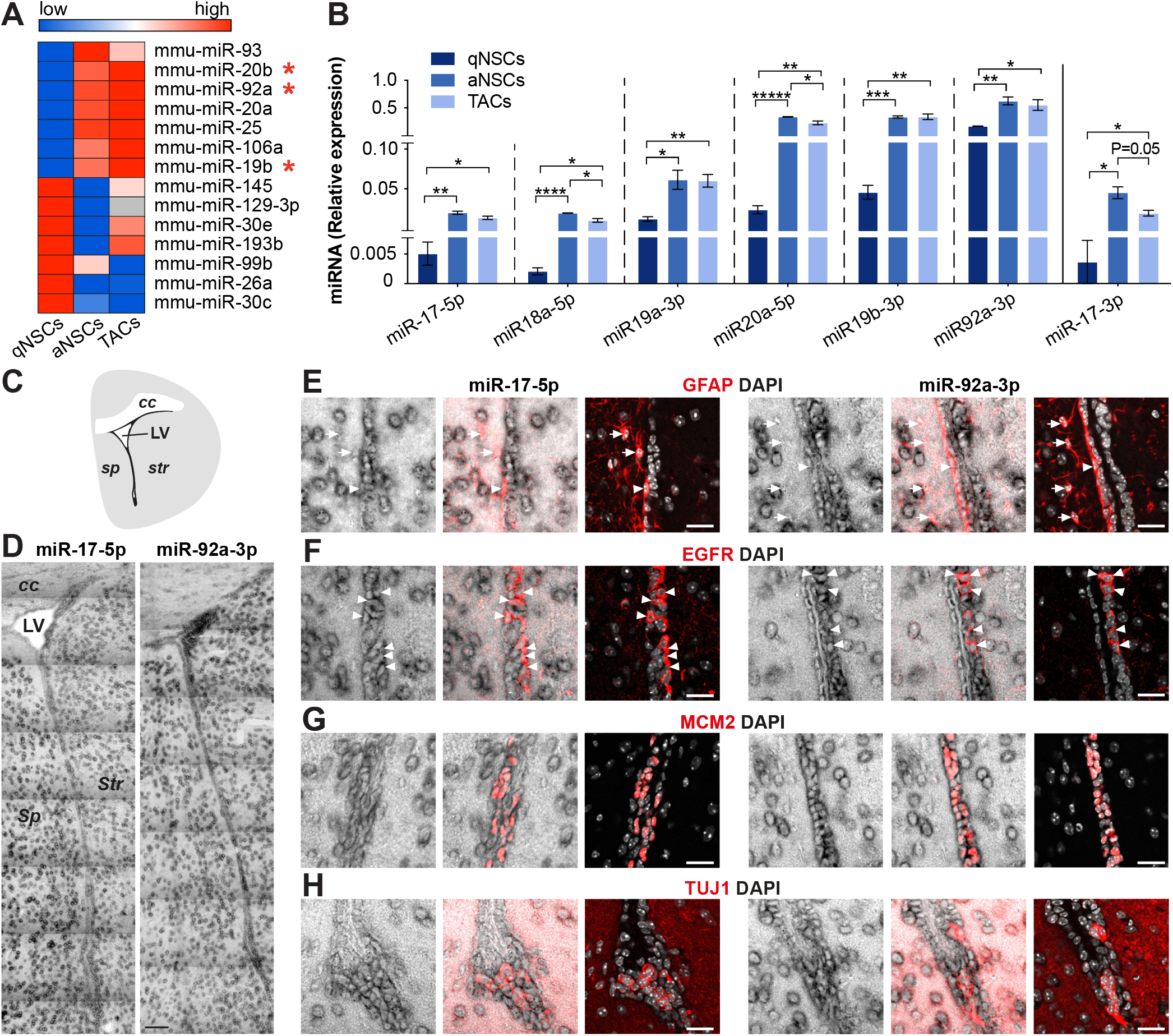
miR-17∼92 is expressed in the adult V-SVZ neurogenic lineage. (A) Differentially expressed miRNAs in FACS-purified qNSCs and aNSCs (unpaired t-test, p < 0.05), and their relative expression in qNSCs, aNSCs and TACs. miRNAs with an asterisk belong to the miR-17∼92 cluster. (B) Relative expression (2^-ΔCt^) of miR-17∼92 members to miR-16-5p in FACS-purified qNSCs, aNSCs and TACs (n = 3; mean ± SEM). (C) Coronal schema of brain showing the lateral ventricle walls and V-SVZ (black). (D) *In situ* hybridization for miR-17-5p and miR-92a-3p in coronal brain sections. (E-H) *In situ* hybridization for miR-17-5p (left) and miR-92a-3p (right) combined with immunostaining in V-SVZ. (E) miR-17-5p and miR-92a-3p were expressed in a few GFAP^+^ (red) radial cells (arrowheads), but not in nearby astrocytes (arrows). miR-17-5p and miR-92a-3p were detected in most EGFR^+^ cells (red, arrowheads) (F), MCM2^+^ proliferating cells (red) (G) and TUJ1^+^ neuroblasts in the V-SVZ (red) (H). Parenchymal mature neurons outside the V-SVZ also expressed miR-17-5p and miR-92a-3p (H). Scale bars: 20um. *cc: corpus callosum, sp: septum, str: striatum*.

To confirm miR-17∼92 expression in the V-SVZ, we performed qPCR analysis for mature miRNAs in FACS-purified cells and *in situ* hybridization (ISH). All guide forms of the miR-17∼92 cluster, as well as the star form miR-17-3p, were expressed at low levels in qNSCs and were significantly upregulated in both aNSCs and TACs (Fig. 1B). ISH revealed similar expression patterns throughout the V-SVZ for different members of the cluster, including miR-17-5p, miR-18-5p, miR-19a-3p and miR-92a-3p (Fig. 1C, D, Suppl. Fig. 1C-E). However, individual miR-17∼92 members showed different expression levels, with miR-92a-3p being the most abundant in the V-SVZ. To define which stages along the V-SVZ lineage express the cluster, we combined immunostaining for lineage markers with ISH. Given the similar distribution of miR-17∼92 miRNAs, we focused on miR-17-5p and miR-92a-3p, the first and the last members of the cluster (Fig. 1E-H, Suppl. Fig. 1B). We detected miR-17-5p and miR-92a-3p in a few GFAP^+^ radial cells (Fig. 1E), consistent with their low level in FACS-purified qNSCs. In contrast, most proliferating cells, including MCM2^+^ cells and EGFR^+^ TACs, had higher levels of miR-17-5p and miR-92a-3p (Fig. 1F, G). TUJ1^+^ neuroblasts also expressed miR-17-5p and miR-92a-3p (Fig. 1H). Outside of the V-SVZ, cluster members were broadly detected throughout the brain (Fig. 1D, Suppl. Fig. 1C-E) in mature parenchymal neurons (Fig. 1H), and rarely in astrocytes (glutamine synthetase^+^ (GS) or GFAP^+^) (Fig. 1E, Suppl. Fig. 1F), as well as in the choroid plexus (Suppl. Fig. 1E). Thus, within the V-SVZ the miR-17∼92 cluster is upregulated upon stem cell activation and is expressed along the neurogenic lineage.

### miR-17∼92 deletion reduces NSC proliferation and neurogenesis

To assess the effect of miR-17∼92 deletion on NSCs *in vitro*, we FACS-purified aNSCs from adult CAGG::CreER^T2+/-^; miR-17∼92^fl/fl^; ROSA^(ACTB-tdTomato,-EGFP)^ (CAGG-miR-17∼92^floxed^) mice. Upon addition of hydroxytamoxifen, the ubiquitously expressed CreER^T2^ recombinase induces deletion of the miR-17∼92 cluster, and reporter expression is switched from tdTomato to eGFP (Fig. 2A). qPCR analysis after cluster deletion *in vitro* confirmed reduced miR-17∼92 levels as assessed by using miR-17-5p as a proxy for the whole cluster (Suppl. Fig. 2A). To define the effect of cluster deletion on stem cell proliferation we used two independent assays, neurospheres and adherent cultures (Fig. 2B). Deletion of the cluster in FACS-purified aNSCs resulted in a 73% reduction in neurosphere formation (Fig. 2C, E). Moreover, deleted neurospheres were visibly smaller than controls and they could not be passaged to form secondary neurospheres (data not shown). In adherent cultures from single FACS-purified aNSCs, miR-17∼92-deleted cells exhibited little proliferation and failed to generate large colonies (>10 cells) as opposed to their control counterparts that could give rise to large clones (>500 cells) (Fig. 2D, F). miR-17∼92 can also play a role in cell survival (Koralov et al., 2008; Olive et al., 2009; Olive et al., 2010; Tung et al., 2015; Ventura et al., 2008; Xiao et al., 2008). However, no differences in apoptotic cells were found between control and deleted cells based on Annexin staining (Suppl. Fig. 2B). Together, these results show an important function of the cluster in adult NSC proliferation *in vitro*, but not in their survival at short time points.

**Figure 2.**
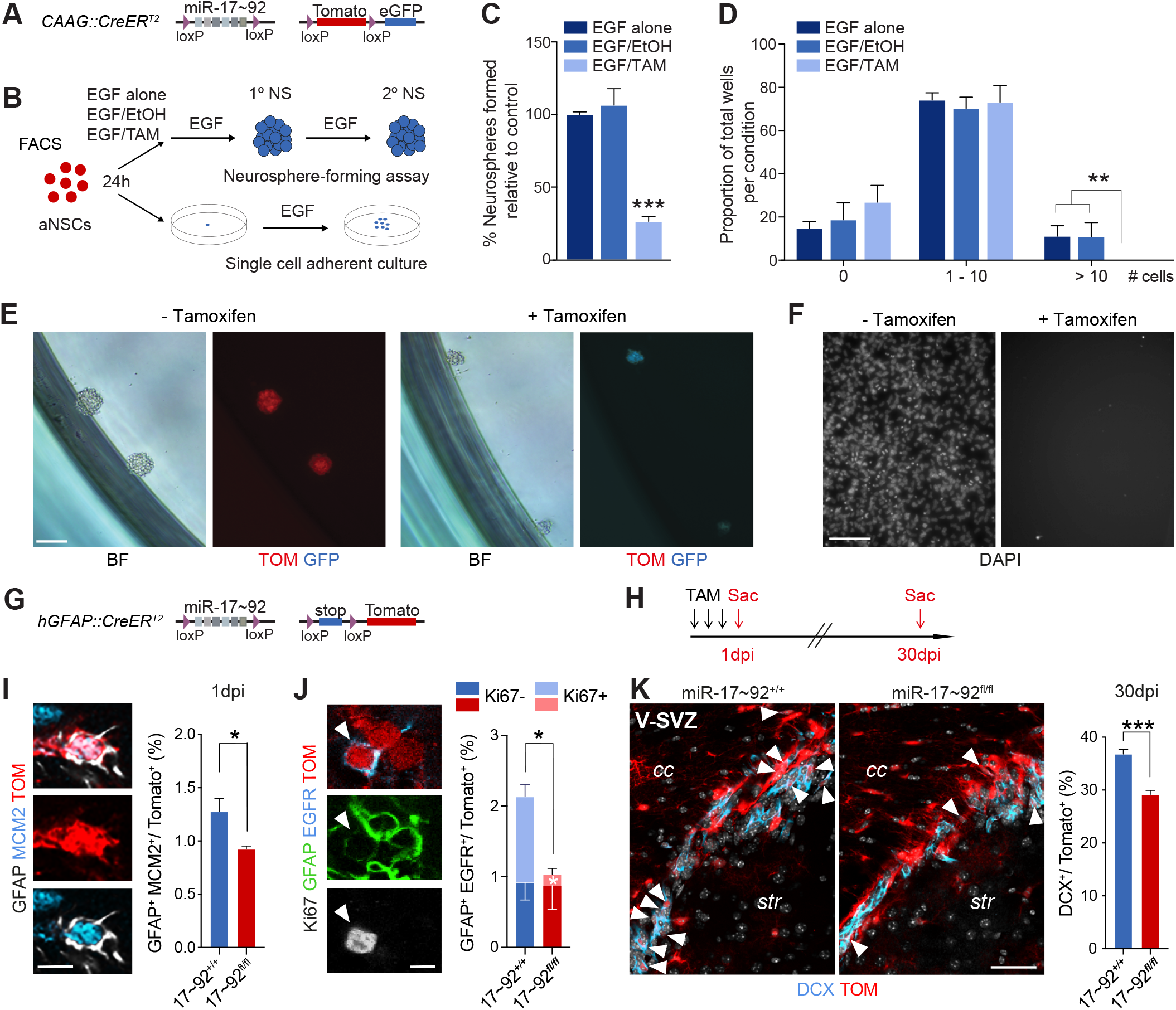
miR-17∼92 deletion in NSCs reduces stem cell proliferation and neurogenesis. (A-F) Deletion of miR-17∼92 decreased NSC proliferation *in vitro*. (A) Schematic of transgenic mouse line used for miR-17∼92 deletion *in vitro*. (B) Schema of experimental paradigms used for *in vitro* analysis. (C, E) miR-17∼92 deletion decreased neurosphere (NS) formation. (C) Quantification of primary neurospheres generated by aNSCs from CAGG-miR-17∼92 ^floxed^ mice at 6 days *in vitro* (div) (n = 3; mean ± SEM). (E) Representative brightfield and confocal images of CAGG::CreERT2^-/-^; miR-17∼92^fl/fl^; ROSA^(ACTB-tdTomato,-EGFP)^ (left) and CAGG-miR-17∼92^floxed^ (right) primary neurospheres at 6 div. Scale bar: 100µm. (D, F) miR-17∼92 deletion decreases proliferation and prevents formation of large colonies. Quantification (D) of total DAPI cells per well in adherent culture after 13 div (n = 3; mean ± SEM). (F) Representative image of large colonies (>500 cells) formed by a single aNSC from CAGG-miR-17∼92 ^floxed^ control cells (EGF/EtOH) (left), but not by deleted cells (right). Scale bar: 100µm. (G-K) Deletion of miR-17∼92 decreased NSC proliferation and neurogenesis *in vivo*. (G) Schematic of transgenic mouse line used for miR-17∼92 deletion *in vivo*. (H) Schema of experimental paradigm used for *in vivo* analysis. (I, J) Confocal images and quantification of TOM^+^ GFAP^+^ MCM2^+^ NSCs (I) and TOM^+^ GFAP^+^ EGFR^+^ Ki67^+/-^ NSCs (J, arrowheads) at 1dpi. (K) Confocal images and quantification of TOM^+^ DCX^+^ neuroblasts in the V-SVZ at 30dpi. (I-K) (n = 3; mean ± SEM). TAM refers to 4-hydroxytamoxifen for *in vitro* and tamoxifen for *in vivo* experiments. Scale bar: (I-J) 5µm, (K) 20µm. cc: *corpus callosum*, str: *striatum*.

To investigate the role of miR-17∼92 in V-SVZ NSCs and their lineage *in vivo*, we selectively deleted the cluster in GFAP^+^ NSCs. To do this, we used adult hGFAP::CreERT2^+/+^; miR-17∼92^+/+^; Ai14 (miR-17∼92^+/+^) and hGFAP::CreER^T2+/+^; miR-17∼92^fl/fl^; Ai14 (miR-17∼92^fl/fl^) mice – in which administration of tamoxifen (TAM) induces miR-17∼92 deletion and initiates tdTomato reporter expression in GFAP^+^ cells (Fig. 2G) – and carried out analyses one and thirty days post TAM injections (1 and 30dpi) (Fig. 2H). qPCR analysis of miR-17-5p, miR-20a-5p and miR-92a-3p in FACS-purified Tomato^+^ (TOM) EGFR^+^ cells (aNSCs and TACs) revealed a 40% to 70% drop in miRNA levels at 1dpi, validating deletion of miR-17∼92 *in vivo* (Suppl Fig. 2C). At 1dpi, consistent with the results from the *in vitro* assays, deletion of the cluster reduced the proportion of dividing NSCs (GFAP^+^ MCM2^+^) (Fig. 2I) and activated NSCs (GFAP^+^ EGFR^+^) (Fig. 2J) over total TOM^+^ cells. The decrease in aNSCs was due to a selective loss of dividing aNSCs (Fig. 2J) indicating that the miR-17∼92 cluster is important for stem cell proliferation *in vivo*. No change was observed in the proportion of total TACs (GFAP^-^ EGFR^+^) (Suppl. Fig. 3A) or dividing TACs (GFAP^-^ EGFR^+^ Ki67^+^) (Suppl. Fig. 3B), neuroblasts (DCX^+^) (Suppl. Fig. 3C) or apoptotic cells (CASP3^+^) (Suppl. Fig. 3D) one day after miR-17∼92 deletion. As astrocytes in tumors release miR-19a-containing exosomes (Zhang et al., 2015), we investigated cell non-autonomous effects of miR-17∼92 deletion in the V-SVZ. Importantly, no changes were detected in Tomato-negative cells, indicating that the effects we observed in the stem cell lineage are cell autonomous (Suppl. Fig. 3F-J). Thirty days after cluster deletion there was an increase in the proportion of TOM^+^ qNSCs paralleled by a decrease in that of TOM^+^ aNSCs in the V-SVZ, consistent with reduced stem cell activation (Suppl. Fig. 2D-F). DCX^+^ neuroblasts were also diminished 30 days after deletion of the cluster (Fig. 2K). Thus, miR-17∼92 deletion in NSCs *in vivo* decreases NSC proliferation and neurogenesis in the V-SVZ.

### miR-17∼92 targets are enriched in stem cell and gliogenesis-related pathways

miRNAs are repressors of gene expression at the post-transcriptional level. The same miRNA can exert different effects in distinct cellular contexts depending on which target genes are expressed. To investigate whether miR-17∼92 might have roles in addition to regulating proliferation in the adult V-SVZ NSC lineage, we performed bioinformatic analysis. As previously reported, the majority of miR-17∼92 targets were observed in gene categories related to ‘cancer’, ‘immune response’, ‘development’ and ‘apoptosis and survival’ (Fig. 3A, Table S2) (Concepcion *et al*., 2012; Han et al., 2015; Olive *et al*., 2010; Ventura *et al*., 2008; Xu et al., 2015). In addition, miR-17∼92 targets were found in pathways upregulated in qNSCs, such as ‘cell adhesion’ and ‘lipid metabolism’ (Baser et al., 2019; Codega *et al*., 2014; Delgado *et al*., 2021; Llorens-Bobadilla et al., 2015), and aNSCs, such as ‘regulation of cell cycle and proliferation’ and ‘stem cell self-renewal and differentiation’. Interestingly, pathway analysis also identified targets related to ‘astrocyte differentiation’ and ‘oligodendrogenesis’, including the master regulator of oligodendrogenesis Olig2, which has been validated as a functional target during spinal cord progenitor patterning in embryogenesis (Chen et al., 2011). In addition to proliferation, miR-17∼92 may therefore be an important regulator of neurogenesis within the adult V-SVZ by repressing gliogenesis.

**Figure 3.**
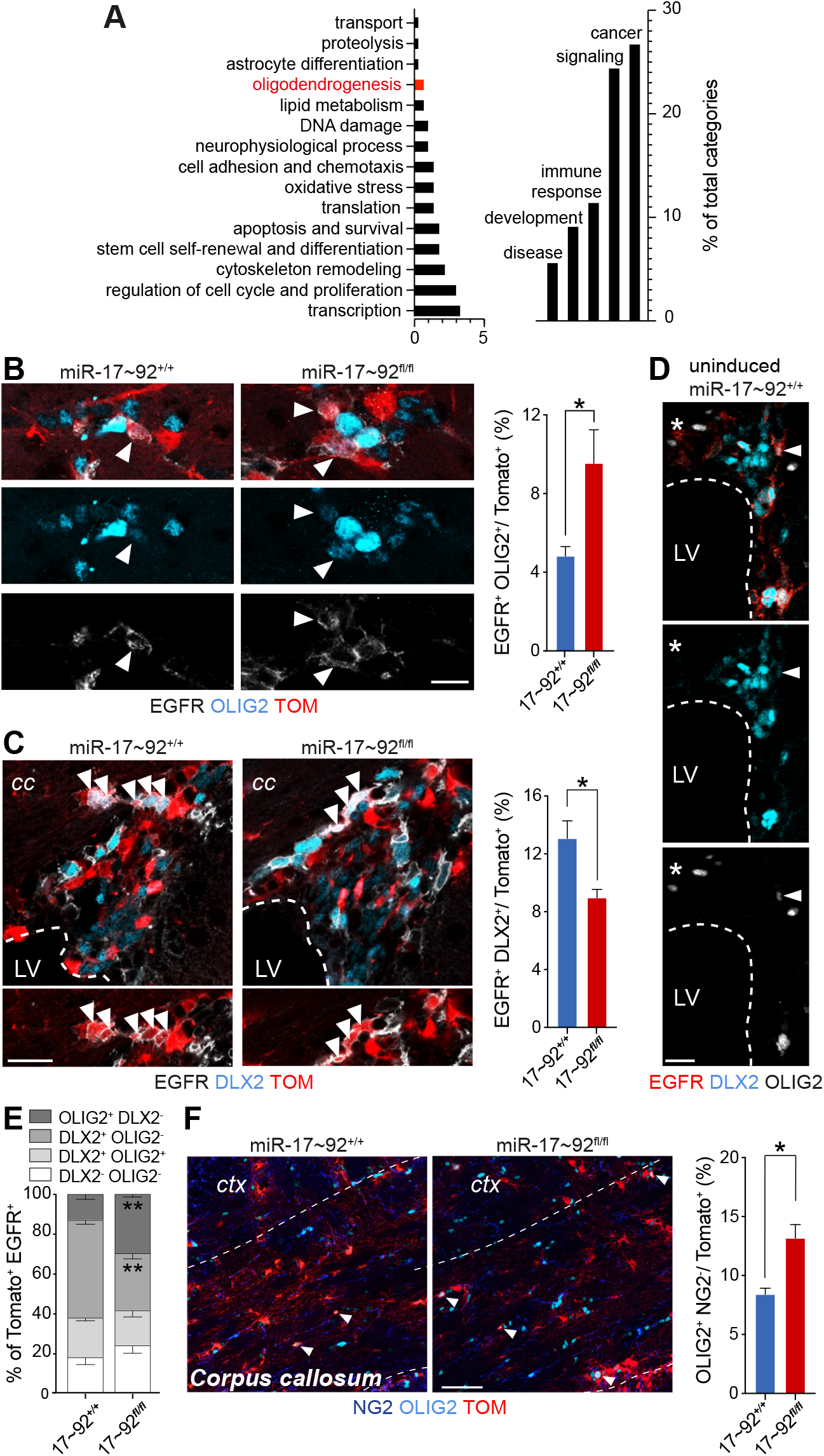
miR-17∼92 deletion leads to a shift from neurogenic to oligodendrogenic transit amplifying cells and increases oligodendrogenesis in *corpus callosum*. (A) Selected top-enriched pathways (MetaCore) containing computationally predicted miR-17∼92 targets expressed in V-SVZ NSCs. Data show pathway abundance as percentage of total categories containing miR-17∼92 targets. (B, C) Confocal images and quantification of TOM^+^ EGFR^+^ OLIG2^+^ TACs (arrowheads) (B) and TOM^+^ EGFR^+^ DLX2^+^ TACs (arrowheads) (C) from miR-17∼92^+/+^ and miR-17∼92^fl/fl^ mice at 1dpi. (D) Confocal micrographs of EGFR (red), DLX2 (blue) and OLIG2 (white) staining on uninduced control miR-17∼92^+/+^ mice. Arrowheads indicate a triple positive cell and asterisks an EGFR^+^ DLX2^-^ OLIG2^-^ cell. (E) Quantification of different TAC (TOM^+^ EGFR^+^) populations in miR-17∼92^+/+^ and miR-17∼92^fl/fl^ mice at 1dpi. (F) Confocal images and quantification of TOM^+^ OLIG2^+^ NG2^-^ cells in the *corpus callosum* from miR-17∼92^+/+^ and miR-17∼92^fl/fl^ mice at 30dpi. (B, C, E, F) n = 3; mean ± SEM. Scale bar: (C) 10µm, (B, D) 20µm, (F) 50µm. cc: *corpus callosum*, ctx: *cortex*, LV: lateral ventricle.

### miR-17∼92 deletion leads to a shift from neurogenic to oligodendrogenic transit amplifying cells and increases oligodendrogenesis in *corpus callosum*

Given the enrichment of miR-17∼92 targets in ‘oligodendrogenesis’, we focused on whether deletion of the cluster may result in a shift towards oligodendrogenesis in the adult V-SVZ. The majority of newly born cells are neuroblasts, generated by DLX2^+^ TACs (Anderson et al., 1997; Brill et al., 2008; Doetsch et al., 2002). However, the V-SVZ also gives rise to oligodendrocytes, via OLIG2^+^ TACs (Menn et al., 2006). Importantly, over-expression and knock-out of Dlx2 or Olig2 is sufficient to shift progenitor cells into a neurogenic or oligodendrogenic fate respectively (Boshans et al., 2021; Jiang et al., 2020; Petryniak et al., 2007).

Since Olig2 is a functional miR-17∼92 target (Chen *et al*., 2011), we postulated that Olig2 expression would increase upon miR-17∼92 deletion. We therefore performed immunostaining for OLIG2 together with the lineage markers GFAP and EGFR at 1dpi. Deletion of the cluster did not affect the proportion of OLIG2^+^ aNSCs (Suppl. Fig. 3E). Importantly, although the proportion of total TACs (Suppl. Fig. 3A) did not change, miR-17∼92 deletion resulted in an increase in TACs expressing OLIG2 (Fig. 3B). Conversely, cluster deletion reduced neurogenic TACs (EGFR^+^ DLX2^+^) (Fig. 3C).

To better understand how TACs were being affected by miR-17∼92 deletion, we first characterized TAC subpopulations in uninduced control mice by performing co-immunostaining for EGFR, DLX2 and OLIG2. This analysis identified four subpopulations of EGFR^+^ TACs. As previously described (Doetsch *et al*., 2002; Menn *et al*., 2006), we detected neurogenic TACs (DLX2^+^ OLIG2^-^) (46% ± 2.5%) and oligodendrogenic TACs (OLIG2^+^ DLX2^-^) (24.5% ± 1%) (Fig. 3D, Suppl. Fig. 3K-L). We also identified a smaller proportion of previously unreported TACs co-expressing DLX2 and OLIG2 (13.5% ± 1.1%), and one negative for both markers (16% ± 2.2%) (Fig. 3D, Suppl. Fig. 3K-L). miR-17∼92 deletion did not change the proportion of OLIG2^+^ DLX2^+^ TACs or double negative TACs (Fig. 3E). Instead, deletion of the cluster led to an increase in the proportion of OLIG2^+^ DLX2^-^ and a decrease in DLX2^+^ OLIG2^-^ TACs (Fig. 3E). Thus, the global increase in OLIG2^+^ TACs was due to a selective shift from neurogenic towards oligodendrogenic TACs. Under normal conditions, the V-SVZ generates a small number of oligodendrocytes that migrate to the *corpus callosum* (Menn *et al*., 2006; Nait-Oumesmar et al., 1999; Ortega et al., 2013). Consistent with the above shift towards oligodendrogenic TACs, mature oligodendrocytes (OLIG2^+^ NG2^-^) were increased in the *corpus callosum* in miR-17∼92 deleted mice at 30 dpi (Fig. 3F). Altogether, these results highlight a role for the miR-17∼92 cluster in repressing gliogenesis.

### miR-17∼92 regulates intraventricular OPC proliferation and maturation

Recently, a population of OPCs located on the ventricular surface of the lateral ventricles (intraventricular OPCs) was described (Delgado *et al*., 2021) (Fig. 4A-B). Intraventricular OPCs are generated within GFAP^+^ clusters (OPCs in clusters) (Fig. 4C). They then disperse from the clusters as immature cells with few processes (immature OPCs) (Fig. 4D), and mature to have more a ramified morphology (mature OPCs) (Fig. 4E). Since deletion of miR-17∼92 affected oligodendrogenesis to the *corpus callosum*, we investigated whether it also affected intraventricular OPCs, using wholemount preparations of miR-17∼92 control and deleted mice (Fig. 4F-M). Consistent with the role of the cluster in proliferation, miR-17∼92 deletion resulted in a reduction in the proportion of intraventricular OPCs expressing MCM2 (Fig. 4I, M, Suppl. Fig. 4A, B). Interestingly, more TOM^+^ OPCs were present in clusters in mutant mice compared to controls (50% versus 15%) (Fig. 4F, G) and more of the clusters were larger (>5 cells) at 1dpi (Fig. 4F, H). The increased cluster size was not due to increased proliferation (Fig. 4I, M, Suppl. Fig. 4A, B) or to defects in survival as assessed by Caspase3 staining (data not shown). Instead, it was likely due to impaired OPC maturation, as the proportion of mature intraventricular OPCs was significantly decreased in mutant mice compared to controls at both at 1dpi and 30dpi (Fig. 4F-G, J-K). Thus, our results indicate a role for the miR-17∼92 cluster in the regulation of intraventricular OPC proliferation and maturation.

**Figure 4.**
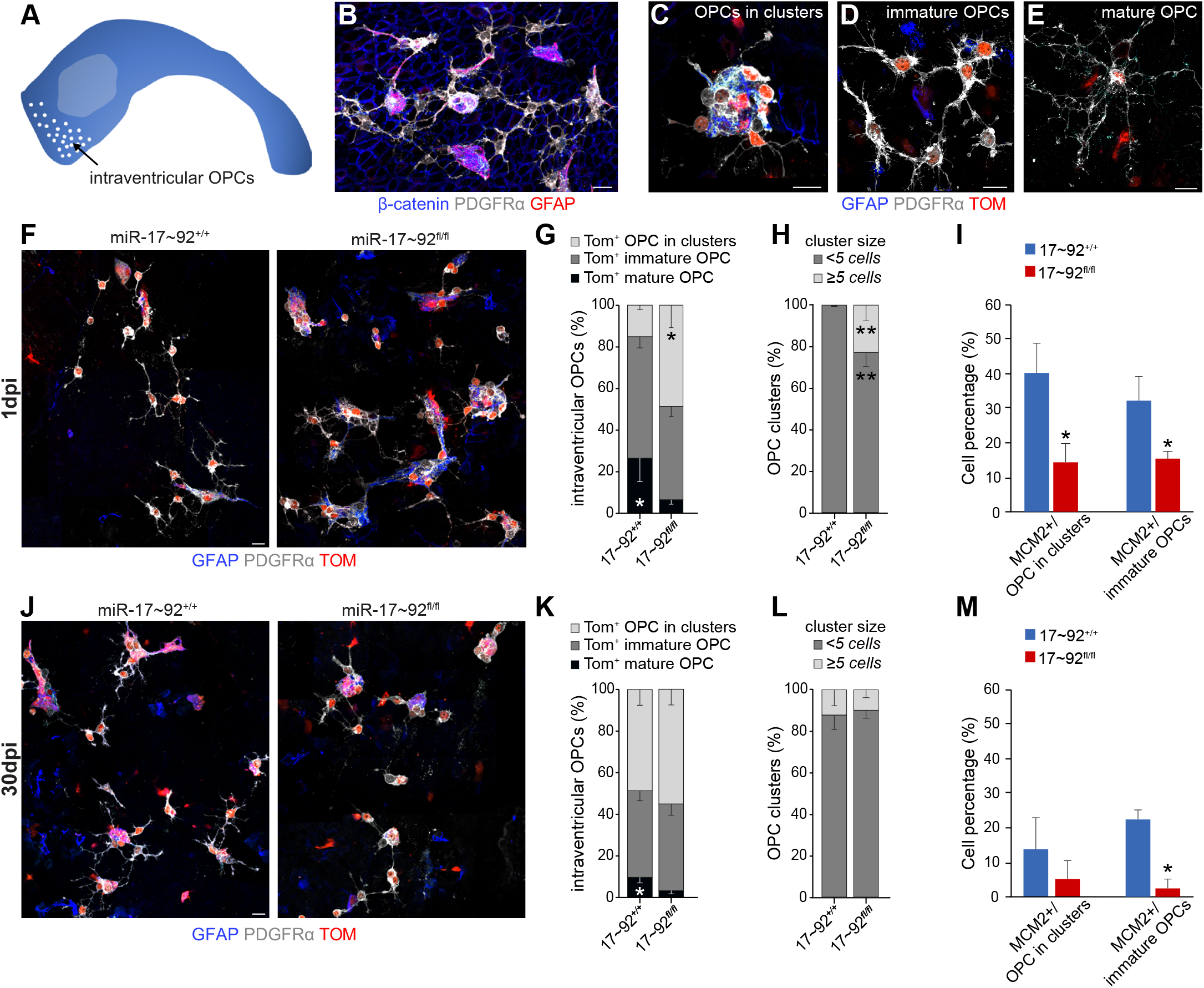
miR-17∼92 regulates intraventricular OPC proliferation and maturation. (A) Schema of wholemount showing location of intraventricular OPCs (white dots). (B-E) Low (B) and high power confocal images (C-E) of PDGFRα^+^ intraventricular OPCs on top of the ventricular surface (β-catenin) at different maturation stages. (F-I) Analysis of intraventricular OPCs in miR-17∼92^+/+^ and miR-17∼92^fl/fl^ at 1dpi. Confocal images of wholemounts (F) and quantification of proportion of intraventricular OPCs at different maturation stages (G), large (≥5 cells) and small (<5 cells) OPC clusters (H), and MCM2^+^ cells at different stages (I). (J-M) Analysis of intraventricular OPCs in miR-17∼92^+/+^ and miR-17∼92^fl/fl^ at 30dpi. Confocal images (J) and quantification of the proportion of intraventricular OPCs at different maturation stages (K), large (≥5 cells) and small (<5 cells) OPC clusters (L) and MCM2^+^ cells at different stages (M). Scale bars: 10µm. (n≥5; mean ± SEM).

## Discussion

miRNAs are emerging as important regulators of adult NSCs. A growing number of miRNAs have been implicated in multiple steps of adult V-SVZ neurogenesis. Among them, miR-106b∼25, miR-137, miR-204 and miR-184 regulate stem cell self-renewal and proliferation (Brett *et al*., 2011; Lepko *et al*., 2019; Liu *et al*., 2010; Szulwach *et al*., 2010), whereas let-7b, miR-7a, miR-9, miR-124 and miR-132 regulate fate specification and functional integration of new neurons into the OB (Akerblom *et al*., 2014; Cheng *et al*., 2009; de Chevigny *et al*., 2012; Pathania *et al*., 2012).

Here, we identify additional miRNA candidates in both qNSCs and aNSCs that might provide further insights into the regulation of early stages of the adult V-SVZ stem cell lineage. Interestingly, all miRNAs enriched in aNSCs compared to qNSCs belonged to the miR-17∼92 family of miRNA clusters, consisting of miR-17∼92, miR-106a∼363 and miR-106b∼25. Notably, although miRNAs from miR-106a∼363 were not previously detected in adult tissues (Ventura *et al*., 2008), our miRNA profiling reveals expression of some miR-106a∼363 members in the adult V-SVZ. While we focused on miR-17∼92, it would be interesting to assess whether miR-106a∼363 also plays a roles in this stem cell niche in the adult.

In the adult V-SVZ, miR-17∼92 is upregulated upon stem cell activation in the neurogenic lineage, and its expression is maintained in TACs and in neuroblasts. In line with its role in cell proliferation and survival in hematopoiesis and in cancers (Mavrakis et al., 2010; Mu et al., 2009; Olive *et al*., 2009; Ventura *et al*., 2008), as well as in adult V-SVZ neural progenitors following experimental stroke (Liu *et al*., 2013), we show that conditional deletion of miR-17∼92 in adult NSCs reduces stem cell proliferation both *in vitro* and *in vivo*. Although we did not observe any defects in survival in miR-17∼92 deleted NSCs at short time points, miR-17∼92 may have a pro-survival function at longer time points or at later stages in the V-SVZ lineage, such as in mature neurons, as was shown during the development of limb innervating motor neurons (Tung *et al*., 2015).

Interestingly, the miR-17∼92 cluster exerts different effects at distinct stages of the lineage, likely depending on the ensemble of targets that are expressed. In TACs, we did not detect an effect on proliferation, but instead observed a shift towards oligodendrogenic TACs, which leads to increased oligodendrocyte formation in the *corpus callosum*. Our data suggest that this shift occurs in TACs themselves, as we did not observe a change in OLIG2^+^ aNSCs. However, as the proportion of total aNSCs is very small following cluster deletion, this shift towards oligodendrogenesis may be initiated earlier in a subset of aNSCs.

miR-17∼92 also acts in the V-SVZ lineage that generates intraventricular OPCs, promoting their proliferation and acquisition of a mature morphology. miR-17∼92 deletion therefore differentially affects V-SVZ oligodendrogenic TACs and intraventricular OPCs, perhaps reflecting different lineages and sets of targets expressed in these two populations. In the adult brain, oligodendrocytes are also generated by parenchymal OPCs, a distinct population widely distributed throughout the brain. It will be interesting to investigate whether adult parenchymal OPC division is also regulated by miR-17∼92, as the cluster positively regulates proliferation in embryonic OPCs (Budde et al., 2010).

NSCs in the adult V-SVZ are heterogeneous at multiple levels, including their developmental origin (Fuentealba et al., 2015; Furutachi et al., 2015; Yuzwa et al., 2017), morphology (Delgado *et al*., 2021; Mirzadeh et al., 2008; Shen et al., 2008), regional position (reviewed in (Azim et al., 2016)) and response to physiological cues (Chaker et al., 2021; Paul et al., 2017). TACs are also heterogeneous with respect to their cell cycle dynamics and transcriptional signatures, which depends on their location (Azim et al., 2015; Ponti et al., 2013). However, to date, the cellular diversity within the TAC population remains poorly explored. Here, we show that four subpopulations of TACs can be distinguished based on the expression of DLX2 and OLIG2: DLX2^-^ OLIG2^-^, DLX2^+^ OLIG2^+^, DLX2^+^ OLIG2^-^ and OLIG2^+^ DLX2^-^, further highlighting the heterogeneity of TACs in the V-SVZ. In the future it will be important to define whether this heterogeneity reflects different stages in the lineage or cells with different potencies, including astrogenic progenitors. Future studies will also be required to clarify whether, in the V-SVZ, fate decisions are made at the stage of NSC or TAC.

miR-17∼92 has been implicated in numerous disorders, including cancer (Concepcion *et al*., 2012; de Pontual et al., 2011; Hayashita et al., 2005; He et al., 2005; Mi et al., 2010; Ota et al., 2004), as well as anxiety and depression (Jin et al., 2016). Moreover, it is upregulated upon stroke in the V-SVZ (Liu *et al*., 2013). In light of our findings, the miR-17∼92 cluster may be downregulated in demyelinating lesions, in which oligodendrogenesis is increased. Altogether, our findings show lineage and cell-type specific effects of the cluster in the adult V-SVZ. As such, regulation of the miR-17∼92 cluster may be an important mechanism to dynamically modulate the generation of distinct neurogenic and gliogenic lineages from adult neural stem cells in response to different physiological and pathological contexts.

## Author Contributions

Conceptualization: F.D., F.F. and A.D. Performed experiments: F.F., A.D. and A.C.D. Data analysis: F.F. and A.D. Supervision: F.D. Manuscript writing: F.D. and F.F.

## Acknowledgements

Work was supported by NIH NINDS R01 NS074039, NIH NINDS R01NS053884, NIH NINDS ARRA NS053884-03S109, NYSTEM C028118, NYSTEM C024287, Swiss National Science Foundation 31003A_163088, European Research Council Advanced Grant (No 789328), and the University of Basel to FD, and NIH T32 GM008224, TL1 TR000082 and NIH NINDS F31NS081990 (A.M.D.). We thank members of the Doetsch and Wichterle labs, J.A. Chen, H. Wichterle, P. Scheiffele and J. Betschinger for discussion; P. Codega, E. Pastrana, V. Crotet and V. Silva-Vargas for help with experiments; J. Zavadil at the Genome Technology Center of NYU Langone Medical Center; the Biozentrum Imaging Core Facility; J. Bögli and S. Stefanova at the Biozentrum FACS Core Facility. We thank J. Chen, H. Wichterle and L. Jeker for mouse lines, T. Sun for sharing the *in situ* hybridization protocol, and C. Schindler for help with statistical analysis.

## Declaration of interests

The authors declare no competing interests.

## Materials and Methods

### Mice

All experiments were performed in accordance with institutional and national guidelines for animal use and approved by IACUC at Columbia University and the cantonal veterinary office of Basel-Stadt. All mice (males and females) were group-housed conventionally with *ad libitum* food and water in 12-h light /12-h dark cycles, and sacrificed at the same time of day. All mice used were three to four months old. The following mouse lines were used: hGFAP::GFP (The Jackson Laboratory, JAX: 003257), CD1(Charles River), RjOrl:SWISS (Janvier Labs), CAGG::CreER^T2^; miR-17∼92^fl/fl^ mice (kindly provided by the Wichterle laboratory at Columbia University), Gt(ROSA)26Sor^tm4(ACTB-tdTomato,-EGFP)Luo^/J mice (ROSA ^(ACTB-tdTomato,-EGFP)^, The Jackson Laboratory, JAX: 007576), miR-17∼92^fl/fl^ mice (miR-17∼92^fl/fl^, The Jackson Laboratory, JAX: 008458, kindly provided by the Jeker laboratory at the University of Basel), B6.Cg-Tg(GFAP-cre/ERT2)505Fmv/J mice (hGFAP::CreER^T2^, The Jackson Laboratory, JAX: 012849) and Gt(ROSA)26Sor^tm14(CAG-tdTomato)Hze^ (Ai14; The Jackson Laboratory, JAX: 007914). We bred CAGG::CreER^T2^; miR-17∼92^fl/fl^ mice to ROSA ^(ACTB-tdTomato,-EGFP)^ mice to generate CAGG::CreER^T2+/-^ or ^-/-^; miR-17∼92^fl/fl^; ROSA^(ACTB-tdTomato,-EGFP)^ mice [CAGG-miR-17∼92^floxed^]. We crossed miR-17∼92^fl/fl^ mice to hGFAP::CreER^T2^ mice and Gt(ROSA)26Sor^tm14(CAG-tdTomato)Hze^ mice to generate hGFAP::CreER^T2^; miR-17∼92^+/+^ or ^fl/fl^; Ai14 mice [miR-17∼92^+/+^ and miR-17∼92^fl/fl^].

### FACS-purification strategy

The V-SVZs were dissected from heterozygous hGFAP::GFP mice (Jackson Labs) or wildtype CD-1 mice (Charles River), digested with papain (Worthington, 1,200 units per 5 mice, 10 min at 37°C) in PIPES solution [120 mM NaCl, 5 mM KCl, 50 mM PIPES (Sigma), 0.6% glucose, 1X Antibiotic/Antimycotic (Gibco), and phenol red (Sigma) in water, pH adjusted to 7.6] and mechanically dissociated to single cells after adding ovomucoid (Worthington, 0.7 mg per 5 mice) and DNAse (Worthington, 1,000 units per 5 mice). Cells were centrifuged for 10 min at 4°C without brake in 22% Percoll (Sigma) to remove myelin. Cell stainings were done in 3 steps: First, cells were incubated for 20 min with PE-conjugated anti-mCD24 (rat monoclonal, 1:500, BD Biosciences, Cat# 553262, RRID:AB_394741) and biotinylated anti-mCD133 (rat monoclonal, 1:300, clone 13A4, Thermo Fisher Scientific, Cat# 13-1331-82, RRID:AB_466591). Cells were washed by centrifugation at 1300rpm for 5min. Next, cells were incubated for 10 min with Streptavidin PE-Cy7 (1:500; Thermo Fisher Scientific, Cat# 25-4317-82, RRID:AB_10116480), and washed by centrifugation. Finally, cells were incubated with A647-complexed EGF (1:300; Molecular Probes, Cat# E-35351) for 15 min, and washed by centrifugation. All stainings and washes were carried out on ice in 1% BSA, 0.1% Glucose HBSS solution. To assess cell viability, 4’,6-diamidino-2-phenylindole (DAPI; 1:1000; Sigma) was added to the cell suspension. All cell populations were isolated in a single sort using a Becton Dickinson FACS Aria II using 13 psi pressure and 100-µm nozzle aperture and were collected in Neurosphere medium (NSM) [DMEM/F12 (Life Technologies) supplemented with 0.6% Glucose (Sigma), 1X Hepes (Life Technologies), 1X Insulin-Selenium-Transferrin (Life Technologies), N-2 (Life Technologies), and B-27 supplement (Life Technologies)]. Gates were set manually using single-color control samples and FMO controls. Data were analyzed with FlowJo 9.3 data analysis software and displayed using bi-exponential scaling.

### miRNA profiling and qPCR analysis

For miRNA profiling, total RNA, including small RNAs, was extracted from FACS-purified populations of qNSCs (hGFAP::GFP^+^, CD133^+^, EGFR^-^, CD24^-^) (n=3 biological replicates), aNSCs (hGFAP::GFP^+^, CD133^+^, EGFR^+^, CD24^-^) (n=3 biological replicates) and TACs (hGFAP::GFP^-^, CD133^-^, EGFR^+^, CD24^-^) (n=2 biological replicates) using the miRNeasy kit (Qiagen). miRNAs were pre-amplified and profiled using TaqMan® Array Rodent MicroRNA A Cards v2.0 A as specified by the manufacturer at the Genome Technology Center of New York University Langone Medical Center. Raw data was loaded into the ExpressionSuite program v1.3 (ThermoFischer) and four control RNAs (U6, sno135, sno202 and miR-16-5p) were used to normalize the dataset.

For qPCR analysis of miRNA expression, miRNA-enriched fractions were extracted from FACS-purified qNSCs, aNSCs and TACs using the miRNeasy kit (Qiagen). cDNA was synthesized using the Exiqon miRCURY LNA Universal RT microRNA kit (Qiagen). qPCR for individual miR-17∼92 members was performed using miRCURY LNA probes and miRCURY LNA SYBR Green PCR Kit (Qiagen).

### In vitro assays

To isolate cells for *in vitro* assays, mixed cohorts of male and female CAGG::CreERT2^+/-^ or ^-/-^; miR-17∼92^fl/fl^; ROSA^(ACTB-tdTomato,-EGFP+/+)^ mice were processed for FACS as described above and stained with CD24-FITC (rat monoclonal, 1:1000; Cat# 553261; RRID:AB_394740), EGF-Alexa647 (1:300; Molecular Probes, Cat# E-35351), and biotinylated anti-mCD133 (rat monoclonal, 1:300, clone 13A4, Thermo Fisher Scientific, Cat# 13-1331-82; RRID:AB_466591), which was followed by secondary staining with PE-Cy7-conjugated streptavidin (1:1000; Thermo Fisher Scientific, Cat# 25-4317-82; RRID:AB_10116480). CD133^+^ EGFR^+^ CD24^-^ cells were FACS-purified to obtain aNSCs and used for neurospheres or single cell adherent assays.

For the neurosphere assay, aNSCs were plated at a density of 100 cells per well in 96-well low-attachment plates (Costar). Cells were grown in Neurosphere medium (NSM, see above) in the presence of 20 ng/ml EGF (Upstate). For the first 24 hours immediately after plating, cells were also treated with 500nM hydroxytamoxifen (4OHT, Sigma) or vehicle alone (ethanol, EtOH). At 24 hours, half of the media was replaced with NSM + EGF to reduce toxicity. The cells were then monitored every 2 days for neurosphere formation and recombination.

Neurospheres were passaged as follows. At 7 days post-plating, the total content of each well was collected and dissociated with 3 mg (600 units) of papain for 10 min at 37°C. The reaction was stopped by adding ovomucoid inhibitor (Worthington, 0.7 mg). DNAse (Worthington, 0.5 mg) was added and cells dissociated to single cells by pipetting. Cells were then plated at a density of 100 cells per well, and monitored for the formation of secondary spheres every two days.

For single cell adherent assays, one cell per well was manually plated into 96 well plates previously coated with Poly-D-Lysine (Sigma, 10 µg/ml) and Fibronectin (Sigma, 2 µg/ml). During the first 24 hours after plating, cells were treated with NSM + EGF, NSM + EGF + 250nM 4OHT, or NSM + EGF + vehicle alone (EtOH). At 24 hours, the entire media was replaced with NSM & EGF to prevent toxicity. Cells were fixed with 4% paraformaldehyde (PFA) in 0.1M phosphate buffer (PB) at 13 days post-plating and stained with DAPI (Sigma). Finally, the total number of DAPI^+^ cells per well was quantified blind to condition.

For survival assays, during the first 24 hours after plating, aNSCs were treated with NSM + EGF, NSM + EGF + 250nM 4OHT, or NSM + EGF + vehicle alone (EtOH). At 24 hours the media was replaced with NSM alone for 5 days. Cells were stained with Annexin V-Alexa647 (1:500, Life Technologies), fixed for 30 minutes with 3.2% PFA, and stained for DAPI (1:1000, Sigma). The wells were imaged on the ZEISS Observer 7.1 at 5X. DAPI^+^ cells were quantified for co-expression with Tomato, eGFP, and Annexin V using the cell counter plugin for FIJI. To assess survival independent of proliferation, each colony was considered as one unit.

To validate deletion of the miR-17∼92 cluster *in vitro*, FACS-purified aNSCs were cultured in NSM + EGF for five days, then the entire media was replaced with NSM + EGF, NSM + EGF + 250nM 4OHT, or NSM + EGF + vehicle alone (EtOH) for two days to induce recombination. Medium was then switched to NSM + EGF for 5 additional days to allow recombined cells to further expand. Cells were collected for sorting by trypsinization for 5 minutes at 37°C, followed by dissociation to single cells by pipetting and passed through a 40-µm filter. DAPI was added prior to FACS to detect dead cells. Cells were gated based on expression of tdTomato (non-recombined cells) and eGFP (recombined cells). RNA was extracted from sorted cells using the Exiqon miRCURY RNA extraction kit, and expression of mature miRs assayed by qPCR using the Exiqon probe for miR-17-5p as described above.

### Tamoxifen injections

Cre-mediated recombination in CreER^T2^ transgenic mice was induced by administration of Tamoxifen (Sigma) dissolved at 30 mg/ml in 90% corn oil with 10% ethanol (Sigma). Mice were intraperitoneally injected at the dose of 120 mg/Kg once per day for three consecutive days and sacrificed one or thirty days after the last tamoxifen injection.

### FACS-purification for validation of miR-17∼92 deletion in vivo

To isolate cells for validating miR-17∼92 deletion *in vivo*, Tomato positive or negative, EGFR^+^ and CD24^-^ cells (corresponding to mixed samples of aNSCs and TACs) were FACS-purified at 1dpi from mixed cohort of male and female miR-17∼92^fl/fl^ mice as described above, using CD24-A700 (rat monoclonal, 1:150, BioLegend, Cat# 101836, RRID:AB_2566730) and EGF-Alexa647 (1:300, Molecular Probes, Cat# E-35351) antibodies. Analysis of miRNA expression levels by qPCR was performed as described above.

### Tissue preparation for in situ hybridization and immunohistochemistry

Mice were anesthetized by intraperitoneal injection of pentobarbital (Esconarkon) and were sacrificed by intracardial perfusion of 4% paraformaldehyde (PFA) in either 0.1M phosphate buffer (PB, for free-floating sections) or 1X phosphate buffer saline (PBS, for cryosections). Brains were extracted from the skull and post-fixed overnight. 25 µm thick free-floating coronal sections were cut using a vibratome (Leica VT1000S). For cryosections, after overnight fixation, brains were cryoprotected by sequential overnight incubation in 15% and 30% sucrose in 1X PBS, snap frozen in -70ºC cold isopentane and stored for at least two days at -80ºC. 12µm thick sections were cut on a cryostat (Leica CM3050S). Wholemounts were dissected as described in (Mirzadeh et al., 2010) from saline-perfused mice and post-fixed overnight in 4% PFA in PB 0.1M solution.

### miRNA in situ hybridization

*In situ* hybridization for miRNA expression was performed on cryosections as described in Jin et al., 2016 using locked nucleic acid (LNA) probes. Briefly, scrambled, miR-17-5p, miR-18a-5p, miR-19a-3p and miR-92a-3p LNA probes (Qiagen) were 3’ end labeled with DIG– ddUTP. After fixation with 4% PFA, acetylation with the acetylation buffer (1.33% Triethanolamine, 0.25% Acetic anhydride, 20 mM HCl), treatment with proteinase K (10 mg/ml, Roche) and pre-hybridization (1X Saline-sodium citrate (SSC) buffer, 50% Formamide, 0.1 mg/ml Salmon Sperm DNA Solution, 1X Denhart, 5 mM EDTA, pH7.5), brain sections were hybridized with DIG-labeled LNA probes at 65ºC overnight. After washing with ice-cold wash buffer (1X SSC, 50% Formamide, 0.1% Tween-20) and 1X Maleic acid buffer containing Tween 20 (MABT), sections were blocked with the blocking buffer (1X MABT, 2% Blocking solution, 20% heat-inactivated sheep serum) and incubated with anti-DIG antibody (1:2’000, Roche) overnight. Brain sections were washed with 1X MABT and the staining buffer (0.1 M NaCl, 50 mM MgCl2, 0.1 M Tris-HCl, pH9.5), stained with NBT/BCIP (Roche) at the room temperature. Finally, sections were washed twice in 1X PBS for 10min at room temperature (RT) and subjected to immunostaining as described below.

### Immunostaining

Wholemounts and tissue sections were incubated in blocking solution [for wholemounts, PBS with 3% bovine serum albumin (BSA) and 1.5% Triton-X100 for whole mounts; for tissue sections, PBS with 10% donkey normal serum supplemented with 0.03% Triton-X100 for antibodies against receptors or 0.3% Triton-X100 for all others] for 1 hour at RT and then incubated in primary antibodies in blocking solution either for 36 hours at 4°C or overnight at RT. After washing, sections were incubated with secondary antibodies for 1 hour at RT. After washing, sections and wholemounts were counterstained with DAPI (Sigma). Sections were mounted on slides with either Aqua Polymount (Brunschwig) or FluorSave™ (Millipore Corporation).

### Antibodies

The following primary antibodies were used: anti-β-Catenin (rabbit polyclonal, 1:200, Cell signaling, Cat# 9587, RRID: AB_10695312); anti-cleaved caspase 3 (rabbit polyclonal, 1:100, Cell Signaling, Cat# 9661, RRID: AB_2341188); anti-DLX2 (mouse polyclonal, 1:50, Santa Cruz, Cat# sc-18140, RRID:AB_2292994); anti-doublecortin, DCX (goat polyclonal, 1:100, Santa Cruz, Cat# sc-8066, RRID: AB_2088494); anti-DCX (guinea pig monoclonal, 1:1000, Millipore, Cat# AB5910, RRID:AB_2230227); anti-DsRed (rabbit polyclonal, 1:500, Clontech, Cat# 632496, RRID: AB_10013483); anti-EGFR (goat polyclonal,1:100, R&D, Cat# AF1280, RRID: AB_354717); anti-EGFR (rabbit monoclonal, 1:100, abcam, Cat# 52894, RRID: AB_869579); anti-GFAP (rat monoclonal, 1:1000, Invitrogen, Cat# 13-0300, RRID:AB_2532994); anti-GFAP (chicken polyclonal, 1:600, Millipore, Cat# PA1-10004, RRID: AB_1074620); anti-Glutamine synthetase (rabbit polyclonal, 1:100, abcam, Cat# ab73593, RRID: AB_2247588); anti-Ki67 (rabbit polyclonal, 1:100, abcam, Cat# ab15580, RRID: AB_443209); anti-Ki67 (rat monoclonal, 1:200, Termo Fisher, Cat# 14-5698-80, RRID:AB_10853185); anti-MCM2 (rabbit monoclonal, 1:1000, Cell signaling, Cat# 3619, RRID: AB_2142137); anti-NG2 (rabbit polyclonal, 1:100, Millipore, Cat# AB5320, RRID: AB_91789); anti-OLIG2 (rabbit polyclonal, 1:100, Millipore, Cat# AB9610, RRID: AB_570666); anti-OLIG2 (goat polyclonal, 1:150, R&D, Cat# AF2418, RRID:AB_2157554); anti-PDGFRα (goat polyclonal, 1:150, R&D, Cat# AF1062, RRID: AB_2236897), anti-TUJ1 (mouse monoclonal, 1:250, Covance, Cat# MMS-435P, RRID:AB_2313773). The following secondary antibodies were used: Alexa Fluor-conjugated (405, 1:250, abcam; 488, 647, 568, 1:600, Molecular probes), Cy3-conjugated (1:1000, Jackson ImmunoResearch).

### Brain section imaging and quantification

Tile scans of the entire dorsoventral extent of the V-SVZ on at least four different rostro-caudal levels (Bregmas: ∼+1mm, ∼+0.70mm, ∼+0.3mm and ∼+0mm) were acquired on a Zeiss LSM800 or LSM880 confocal microscope at 25X or 40X with 1.5µm distance between focal planes. Immunostaining using more than 4 fluorophores was imaged on a NikonTi2 widefield microscope at 40X. Tomato^+^ and Tomato^-^ cells were quantified using the Cell Counter plug-in in FIJI. For each rostro-caudal level, cell percentages of different cell populations over total Tomato^+^ or Tomato^-^ cells were calculated. Cell percentages of the same population from different rostro-caudal levels were averaged and the average values from control and deleted mice were then compared to identify statistically significant differences in cell populations. Images from wholemounts were acquired on Zeiss LSM700 or 800 confocals at 40X. For intraventricular OPCs quantification, at least 4 mice per group were analzyed. Mice with less than 15 recombined intraventricular OPCs per wholemount were excluded.

### Bioinformatic analysis

A list of genes expressed in early stages of the lineage was compiled from gene expression datasets of FACS-purified V-SVZ NSC populations (PDGFRβ^+^ CD133^+^, PDGFRβ^+^ EGFR^+^ and EGFR^+^) (Delgado *et al*., 2021). Computationally predicted targets for the guide and star forms of individual members of the miR-17∼92 cluster (miR-17, 18a, 19a, 20a, 19b, 92a and miR-17*) were generated from the online platforms Targetscan, PicTar, microcosm and miRDB. The gene expression list and miR target list were then compared to find overlap. Finally, the resulting list of miR-17∼92 targets was analzyed using MetaCore (Thompson Reuters, New York, NY) performing enrichment analysis by pathway maps.

### Statistical analysis

For analysis of miRNA microarray data, miRNA expression was first normalized to U6, sno202, sno135, and miR-16-5p, using the same threshold and baseline settings as in (Eriksen et al., 2016; Mestdagh et al., 2014), and differential expression between qNSCs and aNSCs determined by unpaired t-test, (p <0.05). Both normalization and analysis were performed in the ExpressionSuite program v1.3 (ThermoFischer). For qPCR analysis, data were normalized to miR-16-5p expression and analyzed by the 2-ΔΔCT method (Livak & Schmittgen, 2001). For neurosphere experiments, differences in neurosphere formation were assessed using paired two-tailed Student’s t-test using Prism 6 software. For single cell adherent cultures, two-sided Wilcoxon rank sum test was used to assess overall changes in the distribution of the outcomes (given number of cells per well), followed by Fisher’s exact test, to later identify which of the outcomes was responsible for the difference. These analyses were performed using R-commander. For *in vivo* experiments, two-way statistical comparisons were conducted by two-tailed unpaired Student’s test of arcsine-transformed relative values (percentages) using Microsoft Excel. Finally, for pathway analysis, significance was defined by p value < 0.05 and False Discovery Rate < 0.05. For all comparisons, significance was established at *p < 0.05, **p < 0.01, ***p < 0.001, ****p < 0.0001, *****p <0.00001. In all graphs, error bars are standard error of the mean (SEM).

## Data availability

Raw Ct values from miRNA arrays of FACS-purified V-SVZ populations are provided in Table S1.

## Notes

### Competing Interest Statement

The authors have declared no competing interest.

## References

Akerblom, M., Petri, R., Sachdeva, R., Klussendorf, T., Mattsson, B., Gentner, B., and Jakobsson, J. (2014). microRNA-125 distinguishes developmentally generated and adult-born olfactory bulb interneurons. Development 141, 1580–1588. 10.1242/dev.101659.

Anderson, S.A., Qiu, M., Bulfone, A., Eisenstat, D.D., Meneses, J., Pedersen, R., and Rubenstein, J.L. (1997). Mutations of the homeobox genes Dlx-1 and Dlx-2 disrupt the striatal subventricular zone and differentiation of late born striatal neurons. Neuron 19, 27–37. 10.1016/s0896-6273(00)80345-1.

Azim, K., Berninger, B., and Raineteau, O. (2016). Mosaic Subventricular Origins of Forebrain Oligodendrogenesis. Front Neurosci 10, 107. 10.3389/fnins.2016.00107.

Azim, K., Hurtado-Chong, A., Fischer, B., Kumar, N., Zweifel, S., Taylor, V., and Raineteau, O. (2015). Transcriptional Hallmarks of Heterogeneous Neural Stem Cell Niches of the Subventricular Zone. Stem Cells 33, 2232–2242. 10.1002/stem.2017.

Baser, A., Skabkin, M., Kleber, S., Dang, Y., Gülcüler Balta, G.S., Kalamakis, G., Göpferich, M., Ibañez, D.C., Schefzik, R., Lopez, A.S., et al. (2019). Onset of differentiation is post-transcriptionally controlled in adult neural stem cells. Nature 566, 100–104. 10.1038/s41586-019-0888-x.

Bian, S., Hong, J., Li, Q., Schebelle, L., Pollock, A., Knauss, J.L., Garg, V., and Sun, T. (2013). MicroRNA cluster miR-17-92 regulates neural stem cell expansion and transition to intermediate progenitors in the developing mouse neocortex. Cell Rep 3, 1398–1406. 10.1016/j.celrep.2013.03.037.

Boshans, L.L., Soh, H., Wood, W.M., Nolan, T.M., Mandoiu, I.I., Yanagawa, Y., Tzingounis, A.V., and Nishiyama, A. (2021). Direct reprogramming of oligodendrocyte precursor cells into GABAergic inhibitory neurons by a single homeodomain transcription factor Dlx2. Scientific Reports 11, 3552. 10.1038/s41598-021-82931-9.

Brett, J.O., Renault, V.M., Rafalski, V.A., Webb, A.E., and Brunet, A. (2011). The microRNA cluster miR-106b∼25 regulates adult neural stem/progenitor cell proliferation and neuronal differentiation. Aging 3, 108–124. 10.18632/aging.100285.

Brill, M.S., Snapyan, M., Wohlfrom, H., Ninkovic, J., Jawerka, M., Mastick, G.S., Ashery-Padan, R., Saghatelyan, A., Berninger, B., and Götz, M. (2008). A Dlx2- and Pax6-Dependent Transcriptional Code for Periglomerular Neuron Specification in the Adult Olfactory Bulb. The Journal of Neuroscience 28, 6439–6452. 10.1523/jneurosci.0700-08.2008.

Budde, H., Schmitt, S., Fitzner, D., Opitz, L., Salinas-Riester, G., and Simons, M. (2010). Control of oligodendroglial cell number by the miR-17-92 cluster. Development 137, 2127–2132. 10.1242/dev.050633.

Chaker, Z., Codega, P., and Doetsch, F. (2016). A mosaic world: puzzles revealed by adult neural stem cell heterogeneity. Wiley Interdiscip Rev Dev Biol. 10.1002/wdev.248.

Chaker, Z., Segalada, C., and Doetsch, F. (2021). Spatio-Temporal Recruitment of Adult Neural Stem Cells for Transient Neurogenesis During Pregnancy. bioRxiv, 2021.2007.2011.451957. 10.1101/2021.07.11.451957.

Chen, J.A., Huang, Y.P., Mazzoni, E.O., Tan, G.C., Zavadil, J., and Wichterle, H. (2011). Mir-17-3p controls spinal neural progenitor patterning by regulating Olig2/Irx3 cross-repressive loop. Neuron 69, 721–735. 10.1016/j.neuron.2011.01.014.

Cheng, L.C., Pastrana, E., Tavazoie, M., and Doetsch, F. (2009). miR-124 regulates adult neurogenesis in the subventricular zone stem cell niche. Nat Neurosci 12, 399–408. 10.1038/nn.2294.

Codega, P., Silva-Vargas, V., Paul, A., Maldonado-Soto, A.R., Deleo, A.M., Pastrana, E., and Doetsch, F. (2014). Prospective identification and purification of quiescent adult neural stem cells from their in vivo niche. Neuron 82, 545–559. 10.1016/j.neuron.2014.02.039.

Concepcion, C.P., Bonetti, C., and Ventura, A. (2012). The microRNA-17-92 family of microRNA clusters in development and disease. Cancer J 18, 262–267. 10.1097/PPO.0b013e318258b60a.

de Chevigny, A., Core, N., Follert, P., Gaudin, M., Barbry, P., Beclin, C., and Cremer, H. (2012). miR-7a regulation of Pax6 controls spatial origin of forebrain dopaminergic neurons. Nat Neurosci 15, 1120–1126. 10.1038/nn.3142.

de Pontual, L., Yao, E., Callier, P., Faivre, L., Drouin, V., Cariou, S., Van Haeringen, A., Geneviève, D., Goldenberg, A., Oufadem, M., et al. (2011). Germline deletion of the miR-17∼92 cluster causes skeletal and growth defects in humans. Nat Genet 43, 1026–1030. 10.1038/ng.915.

Delgado, A.C., Maldonado-Soto, A.R., Silva-Vargas, V., Mizrak, D., von Känel, T., Tan, K.R., Paul, A., Madar, A., Cuervo, H., Kitajewski, J., et al. (2021). Release of stem cells from quiescence reveals gliogenic domains in the adult mouse brain. Science 372, 1205–1209. 10.1126/science.abg8467.

Doetsch, F., Petreanu, L., Caille, I., Garcia-Verdugo, J.-M., and Alvarez-Buylla, A. (2002). EGF Converts Transit-Amplifying Neurogenic Precursors in the Adult Brain into Multipotent Stem Cells. Neuron 36, 1021-1034. https://doi.org/10.1016/S0896-6273(02)01133-9.

Eriksen, A.H., Andersen, R.F., Pallisgaard, N., Sorensen, F.B., Jakobsen, A., and Hansen, T.F. (2016). MicroRNA Expression Profiling to Identify and Validate Reference Genes for the Relative Quantification of microRNA in Rectal Cancer. PLoS One 11, e0150593. 10.1371/journal.pone.0150593.

Fuentealba, L.C., Rompani, S.B., Parraguez, J.I., Obernier, K., Romero, R., Cepko, C.L., and Alvarez-Buylla, A. (2015). Embryonic Origin of Postnatal Neural Stem Cells. Cell 161, 1644–1655. 10.1016/j.cell.2015.05.041.

Furutachi, S., Miya, H., Watanabe, T., Kawai, H., Yamasaki, N., Harada, Y., Imayoshi, I., Nelson, M., Nakayama, K.I., Hirabayashi, Y., and Gotoh, Y. (2015). Slowly dividing neural progenitors are an embryonic origin of adult neural stem cells. Nat Neurosci 18, 657–665. 10.1038/nn.3989.

Han, Y.C., Vidigal, J.A., Mu, P., Yao, E., Singh, I., Gonzalez, A.J., Concepcion, C.P., Bonetti, C., Ogrodowski, P., Carver, B., et al. (2015). An allelic series of miR-17 approximately 92-mutant mice uncovers functional specialization and cooperation among members of a microRNA polycistron. Nat Genet 47, 766–775. 10.1038/ng.3321.

Hayashita, Y., Osada, H., Tatematsu, Y., Yamada, H., Yanagisawa, K., Tomida, S., Yatabe, Y., Kawahara, K., Sekido, Y., and Takahashi, T. (2005). A polycistronic microRNA cluster, miR-17-92, is overexpressed in human lung cancers and enhances cell proliferation. Cancer Res 65, 9628–9632. 10.1158/0008-5472.Can-05-2352.

He, L., Thomson, J.M., Hemann, M.T., Hernando-Monge, E., Mu, D., Goodson, S., Powers, S., Cordon-Cardo, C., Lowe, S.W., Hannon, G.J., and Hammond, S.M. (2005). A microRNA polycistron as a potential human oncogene. Nature 435, 828–833. 10.1038/nature03552.

Jiang, Q., Zagozewski, J., Godbout, R., and Eisenstat, D.D. (2020). Distal-less genes <em>Dlx1/Dlx2</em> repress oligodendrocyte genesis through transcriptional inhibition of <em>Olig2</em> expression in the developing vertebrate forebrain. bioRxiv, 2020.2004.2009.012385. 10.1101/2020.04.09.012385.

Jin, J., Kim, S.N., Liu, X., Zhang, H., Zhang, C., Seo, J.S., Kim, Y., and Sun, T. (2016). miR-17-92 Cluster Regulates Adult Hippocampal Neurogenesis, Anxiety, and Depression. Cell Rep 16, 1653–1663. 10.1016/j.celrep.2016.06.101.

Koralov, S.B., Muljo, S.A., Galler, G.R., Krek, A., Chakraborty, T., Kanellopoulou, C., Jensen, K., Cobb, B.S., Merkenschlager, M., Rajewsky, N., and Rajewsky, K. (2008). Dicer ablation affects antibody diversity and cell survival in the B lymphocyte lineage. Cell 132, 860–874. 10.1016/j.cell.2008.02.020.

Lepko, T., Pusch, M., Müller, T., Schulte, D., Ehses, J., Kiebler, M., Hasler, J., Huttner, H.B., Vandenbroucke, R.E., Vandendriessche, C., et al. (2019). Choroid plexus-derived miR-204 regulates the number of quiescent neural stem cells in the adult brain. The EMBO Journal 38. 10.15252/embj.2018100481.

Liu, C., Teng, Z.Q., Santistevan, N.J., Szulwach, K.E., Guo, W., Jin, P., and Zhao, X. (2010). Epigenetic regulation of miR-184 by MBD1 governs neural stem cell proliferation and differentiation. Cell Stem Cell 6, 433–444. 10.1016/j.stem.2010.02.017.

Liu, X.S., Chopp, M., Wang, X.L., Zhang, L., Hozeska-Solgot, A., Tang, T., Kassis, H., Zhang, R.L., Chen, C., Xu, J., and Zhang, Z.G. (2013). MicroRNA-17-92 cluster mediates the proliferation and survival of neural progenitor cells after stroke. J Biol Chem 288, 12478–12488. 10.1074/jbc.M112.449025.

Llorens-Bobadilla, E., Zhao, S., Baser, A., Saiz-Castro, G., Zwadlo, K., and Martin-Villalba, A. (2015). Single-Cell Transcriptomics Reveals a Population of Dormant Neural Stem Cells that Become Activated upon Brain Injury. Cell Stem Cell 17, 329–340. 10.1016/j.stem.2015.07.002.

Mavrakis, K.J., Wolfe, A.L., Oricchio, E., Palomero, T., de Keersmaecker, K., McJunkin, K., Zuber, J., James, T., Khan, A.A., Leslie, C.S., et al. (2010). Genome-wide RNA-mediated interference screen identifies miR-19 targets in Notch-induced T-cell acute lymphoblastic leukaemia. Nat Cell Biol 12, 372–379. 10.1038/ncb2037.

Menn, B., Garcia-Verdugo, J.M., Yaschine, C., Gonzalez-Perez, O., Rowitch, D., and Alvarez-Buylla, A. (2006). Origin of oligodendrocytes in the subventricular zone of the adult brain. J Neurosci 26, 7907–7918. 10.1523/JNEUROSCI.1299-06.2006.

Mestdagh, P., Hartmann, N., Baeriswyl, L., Andreasen, D., Bernard, N., Chen, C., Cheo, D., D’Andrade, P., DeMayo, M., Dennis, L., et al. (2014). Evaluation of quantitative miRNA expression platforms in the microRNA quality control (miRQC) study. Nature Methods 11, 809–815. 10.1038/nmeth.3014.

Mi, S., Li, Z., Chen, P., He, C., Cao, D., Elkahloun, A., Lu, J., Pelloso, L.A., Wunderlich, M., Huang, H., et al. (2010). Aberrant overexpression and function of the miR-17-92 cluster in MLL-rearranged acute leukemia. Proc Natl Acad Sci U S A 107, 3710–3715. 10.1073/pnas.0914900107.

Mirzadeh, Z., Doetsch, F., Sawamoto, K., Wichterle, H., and Alvarez-Buylla, A. (2010). The Subventricular Zone En-face: Wholemount Staining and Ependymal Flow. JoVE, e1938. doi:10.3791/1938.

Mirzadeh, Z., Merkle, F.T., Soriano-Navarro, M., Garcia-Verdugo, J.M., and Alvarez-Buylla, A. (2008). Neural stem cells confer unique pinwheel architecture to the ventricular surface in neurogenic regions of the adult brain. Cell Stem Cell 3, 265–278. 10.1016/j.stem.2008.07.004.

Mu, P., Han, Y.C., Betel, D., Yao, E., Squatrito, M., Ogrodowski, P., de Stanchina, E., D’Andrea, A., Sander, C., and Ventura, A. (2009). Genetic dissection of the miR-17∼92 cluster of microRNAs in Myc-induced B-cell lymphomas. Genes Dev 23, 2806–2811. 10.1101/gad.1872909.

Nait-Oumesmar, B., Decker, L., Lachapelle, F., Avellana-Adalid, V., Bachelin, C., and Van Evercooren, A.B.-. (1999). Progenitor cells of the adult mouse subventricular zone proliferate, migrate and differentiate into oligodendrocytes after demyelination. European Journal of Neuroscience 11, 4357–4366. https://doi.org/10.1046/j.1460-9568.1999.00873.x.

Naka-Kaneda, H., Nakamura, S., Igarashi, M., Aoi, H., Kanki, H., Tsuyama, J., Tsutsumi, S., Aburatani, H., Shimazaki, T., and Okano, H. (2014). The miR-17/106-p38 axis is a key regulator of the neurogenic-to-gliogenic transition in developing neural stem/progenitor cells. Proc Natl Acad Sci U S A 111, 1604–1609. 10.1073/pnas.1315567111.

O’Brien, J., Hayder, H., Zayed, Y., and Peng, C. (2018). Overview of MicroRNA Biogenesis, Mechanisms of Actions, and Circulation. Front Endocrinol (Lausanne) 9, 402. 10.3389/fendo.2018.00402.

Obernier, K., and Alvarez-Buylla, A. (2019). Neural stem cells: origin, heterogeneity and regulation in the adult mammalian brain. Development 146. 10.1242/dev.156059.

Olive, V., Bennett, M.J., Walker, J.C., Ma, C., Jiang, I., Cordon-Cardo, C., Li, Q.J., Lowe, S.W., Hannon, G.J., and He, L. (2009). miR-19 is a key oncogenic component of mir-17-92. Genes Dev 23, 2839–2849. 10.1101/gad.1861409.

Olive, V., Jiang, I., and He, L. (2010). mir-17-92, a cluster of miRNAs in the midst of the cancer network. Int J Biochem Cell Biol 42, 1348–1354. 10.1016/j.biocel.2010.03.004.

Ortega, F., Gascon, S., Masserdotti, G., Deshpande, A., Simon, C., Fischer, J., Dimou, L., Chichung Lie, D., Schroeder, T., and Berninger, B. (2013). Oligodendrogliogenic and neurogenic adult subependymal zone neural stem cells constitute distinct lineages and exhibit differential responsiveness to Wnt signalling. Nat Cell Biol 15, 602–613. 10.1038/ncb2736.

Ota, A., Tagawa, H., Karnan, S., Tsuzuki, S., Karpas, A., Kira, S., Yoshida, Y., and Seto, M. (2004). Identification and characterization of a novel gene, C13orf25, as a target for 13q31-q32 amplification in malignant lymphoma. Cancer Res 64, 3087–3095. 10.1158/0008-5472.can-03-3773.

Pastrana, E., Cheng, L.C., and Doetsch, F. (2009). Simultaneous prospective purification of adult subventricular zone neural stem cells and their progeny. Proc Natl Acad Sci U S A 106, 6387–6392. 10.1073/pnas.0810407106.

Pathania, M., Torres-Reveron, J., Yan, L., Kimura, T., Lin, T.V., Gordon, V., Teng, Z.Q., Zhao, X., Fulga, T.A., Van Vactor, D., and Bordey, A. (2012). miR-132 enhances dendritic morphogenesis, spine density, synaptic integration, and survival of newborn olfactory bulb neurons. PLoS One 7, e38174. 10.1371/journal.pone.0038174.

Paul, A., Chaker, Z., and Doetsch, F. (2017). Hypothalamic regulation of regionally distinct adult neural stem cells and neurogenesis. Science 356, 1383–1386. 10.1126/science.aal3839.

Petryniak, M.A., Potter, G.B., Rowitch, D.H., and Rubenstein, J.L. (2007). Dlx1 and Dlx2 control neuronal versus oligodendroglial cell fate acquisition in the developing forebrain. Neuron 55, 417–433. 10.1016/j.neuron.2007.06.036.

Ponti, G., Obernier, K., Guinto, C., Jose, L., Bonfanti, L., and Alvarez-Buylla, A. (2013). Cell cycle and lineage progression of neural progenitors in the ventricular-subventricular zones of adult mice. Proceedings of the National Academy of Sciences 110, E1045–E1054. 10.1073/pnas.1219563110.

Shen, Q., Wang, Y., Kokovay, E., Lin, G., Chuang, S.M., Goderie, S.K., Roysam, B., and Temple, S. (2008). Adult SVZ stem cells lie in a vascular niche: a quantitative analysis of niche cell-cell interactions. Cell Stem Cell 3, 289–300. 10.1016/j.stem.2008.07.026.

Szulwach, K.E., Li, X., Smrt, R.D., Li, Y., Luo, Y., Lin, L., Santistevan, N.J., Li, W., Zhao, X., and Jin, P. (2010). Cross talk between microRNA and epigenetic regulation in adult neurogenesis. J Cell Biol 189, 127–141. 10.1083/jcb.200908151.

Tung, Y.T., Lu, Y.L., Peng, K.C., Yen, Y.P., Chang, M., Li, J., Jung, H., Thams, S., Huang, Y.P., Hung, J.H., and Chen, J.A. (2015). Mir-17 approximately 92 Governs Motor Neuron Subtype Survival by Mediating Nuclear PTEN. Cell Rep 11, 1305–1318. 10.1016/j.celrep.2015.04.050.

Ventura, A., Young, A.G., Winslow, M.M., Lintault, L., Meissner, A., Erkeland, S.J., Newman, J., Bronson, R.T., Crowley, D., Stone, J.R., et al. (2008). Targeted deletion reveals essential and overlapping functions of the miR-17 through 92 family of miRNA clusters. Cell 132, 875–886. 10.1016/j.cell.2008.02.019.

Xiao, C., Srinivasan, L., Calado, D.P., Patterson, H.C., Zhang, B., Wang, J., Henderson, J.M., Kutok, J.L., and Rajewsky, K. (2008). Lymphoproliferative disease and autoimmunity in mice with increased miR-17-92 expression in lymphocytes. Nat Immunol 9, 405–414. 10.1038/ni1575.

Xu, S., Ou, X., Huo, J., Lim, K., Huang, Y., Chee, S., and Lam, K.P. (2015). Mir-17-92 regulates bone marrow homing of plasma cells and production of immunoglobulin G2c. Nat Commun 6, 6764. 10.1038/ncomms7764.

Yuzwa, S.A., Borrett, M.J., Innes, B.T., Voronova, A., Ketela, T., Kaplan, D.R., Bader, G.D., and Miller, F.D. (2017). Developmental Emergence of Adult Neural Stem Cells as Revealed by Single-Cell Transcriptional Profiling. Cell Rep 21, 3970–3986. 10.1016/j.celrep.2017.12.017.

Zhang, L., Zhang, S., Yao, J., Lowery, F.J., Zhang, Q., Huang, W.C., Li, P., Li, M., Wang, X., Zhang, C., et al. (2015). Microenvironment-induced PTEN loss by exosomal microRNA primes brain metastasis outgrowth. Nature 527, 100–104. 10.1038/nature15376.

Zhao, C., Sun, G., Li, S., Lang, M.F., Yang, S., Li, W., and Shi, Y. (2010). MicroRNA let-7b regulates neural stem cell proliferation and differentiation by targeting nuclear receptor TLX signaling. Proc Natl Acad Sci U S A 107, 1876–1881. 10.1073/pnas.0908750107.

Zhao, C., Sun, G., Li, S., and Shi, Y. (2009). A feedback regulatory loop involving microRNA-9 and nuclear receptor TLX in neural stem cell fate determination. Nat Struct Mol Biol 16, 365–371. 10.1038/nsmb.1576.

