## Supplementary Information for "miR-17∼92 exerts stage-specific effects in adult V-SVZ neural stem cell lineages"

Includes:

Supplementary Figures and Legends

Supplementary Table Legends

**Supplementary Figure 1. miR-17~92 expression in the adult V-SVZ.**

(A) Schema of coronal section of mouse brain showing the V-SVZ (red) next to the lateral ventricle (LV) and of V-SVZ early lineage and markers for FACS-purification. (B) Schematic representation of the miR-17~92 family of miRNA clusters. miRNAs with homologous seed sequences are in the same color. (C-E) *In situ* hybridization for scrambled, miR-17-5p, miR-18a-5p, miR-19a-3p and miR-92a-3p on V-SVZ coronal sections at different rostrocaudal levels (Bregma +1.4mm (C), +0.7mm (D) and -0.1mm (E)). Scale bars: 50µm. (F) Confocal micrographs of glutamine synthetase$^{+}$ (GS) striatal astrocytes (red) expressing miR-17-5p (arrowheads) (left), miR-19a-3p (arrowheads) (middle) and miR-92a-3p (arrowheads) (right). Scale bar: 20µm. CP: *choroid plexus*, *cc: corpus callosum; sp: septum*, *str: striatum.*

**Supplementary Figure 2. Validation of miR-17~92 deletion and effect on NSC survival and proliferation.**

(A) Schema of experimental paradigm used to validate deletion of miR-17-5p *in vitro* (left). qPCR analysis for miR-17-5p in FACS-purified recombined and non-recombined cells after *in vitro* deletion of the cluster (right) (n=2; mean ± SEM). (B) Schema of experimental paradigm used to assess cell survival *in vitro* (left). Quantification of proportion of Annexin$^{+}$ cells (right) in non-recombined and recombined cells (n = 3; mean ± SEM). (C) Schema of experimental paradigm used to validate *in vivo* deletion in miR-17~92$^{fl/fl}$ mice by qPCR analysis of FACS-purified cells for miR-17-5p, miR-20a-5p and miR-92a-3p (n = 3; mean ± SEM). (D-F) *In vivo* analysis of NSC proliferation at 30dpi. (D) Confocal images of V-SVZ coronal sections and quantification of Tom$^{+}$ qNSCs (E) and aNSCs (F) from miR-17~92$^{+/+}$ and miR-17~92$^{fl/fl}$ mice (n = 3; mean ± SEM). TAM refers to 4-hydroxytamoxifen for *in vitro* and tamoxifen for *in vivo* experiments. Scale bar: 10µm. n.s. non-significant.

**Supplementary Figure 3. *In vivo* analysis of V-SVZ populations in miR-17~92$^{+/+}$ and miR-17~92$^{fl/fl}$ mice.**

(A-E) *In vivo* analysis of Tomato$^{+}$ V-SVZ populations in coronal sections from miR-17~92$^{+/+}$ and miR-17~92$^{fl/fl}$ at 1dpi. Quantification of Tomato$^{+}$ cells in the V-SVZ revealed no differences in (A) total TACs, (B) Ki67$^{+}$ dividing TACs, (C) DCX$^{+}$ neuroblasts, (D) CASP3$^{+}$ apoptotic cells and (E) GFAP$^{+}$ EGFR$^{+}$ OLIG2$^{+}$ aNSCs. (F-J) Quantification of Tomato$^{-}$ cells in the V-SVZ of miR-17~92$^{+/+}$ and miR-17~92$^{fl/fl}$ mice at 1dpi showed no changes in (F) GFAP$^{+}$ EGFR$^{-}$ cells (qNSCs and astrocytes), (G) GFAP$^{+}$ EGFR$^{+}$ aNSCs, (H) GFAP$^{-}$ EGFR$^{+}$ TACs, (I) DCX$^{+}$ neuroblasts and (J) OLIG2$^{+}$ cells (OPCs and oligodendrocytes). (K) Schema of the V-SVZ showing the location of confocal images in K' and K". (K'-K") Confocal images of V-SVZ coronal sections from uninduced miR-17~92$^{+/+}$ mice immunostained for EGFR (magenta), DLX2 (green) and OLIG2 (white). Arrowheads indicate triple positive cells and asterisks an EGFR$^{+}$ DLX2$^{-}$ OLIG2$^{-}$ cell. (L) Quantification of EGFR/DLX2/OLIG2 staining in uninduced miR-17~92$^{+/+}$ mice. Scale bar: 10µm. (A-J, L) n = 3; mean ± SEM; n.s. non-significant.

**Supplementary Figure 4. miR-17~92 decreases intraventricular OPC proliferation.**
Confocal images of wholemounts of the lateral wall showing intraventricular OPCs immunostained for PDGFRα and MCM2 in miR-17~92$^{+/+}$ and miR-17~92$^{fl/fl}$ mice at 1dpi (A) and 30dpi (B). Scale bar: 20µm.

**Table S1. miRNA profiling of FACS-purified qNSCs, aNSCs and TACs**
Ct values for miRNA profiling and differentially expressed miRNAs between qNSCs and aNSCs

**Table S2. Pathway maps containing computationally predicted miR-17~92 targets expressed in V-SVZ NSCs**

### *Supplementary Figure 1*

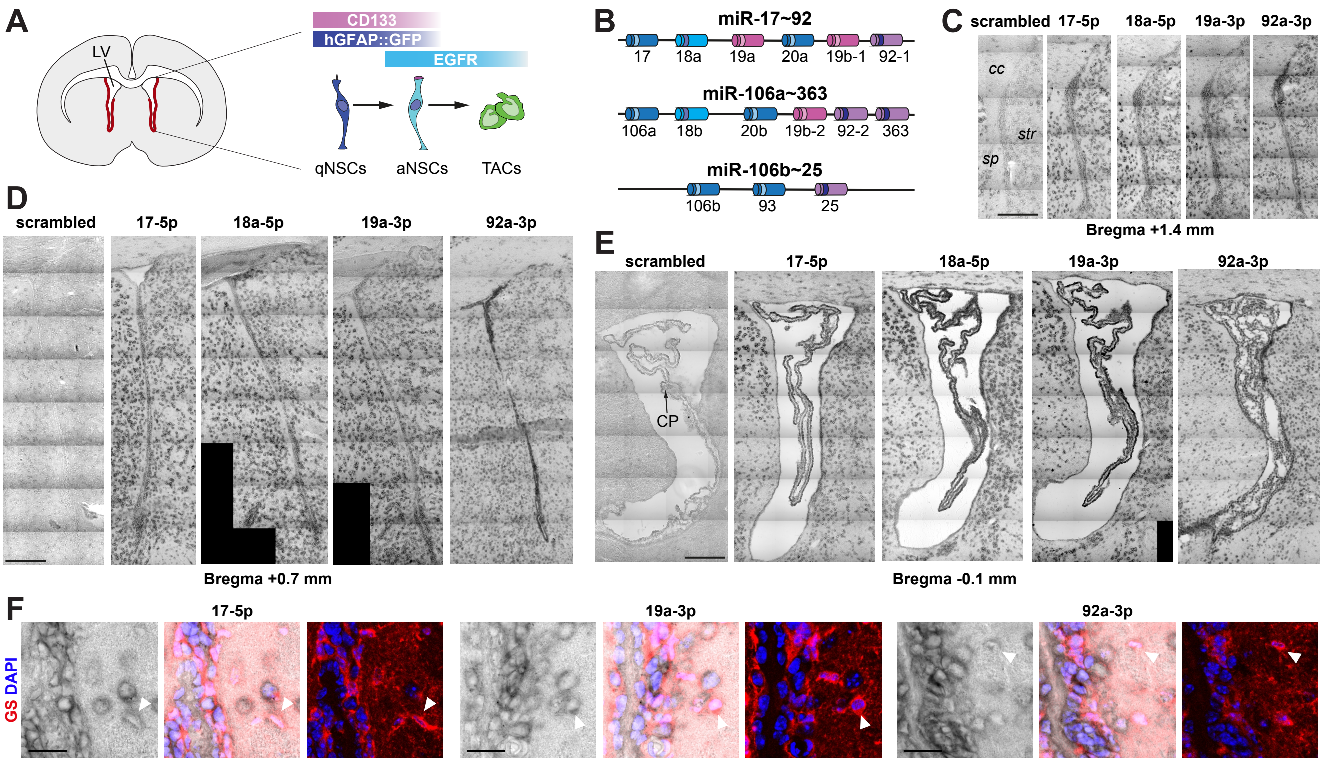

### *Supplementary Figure 2*

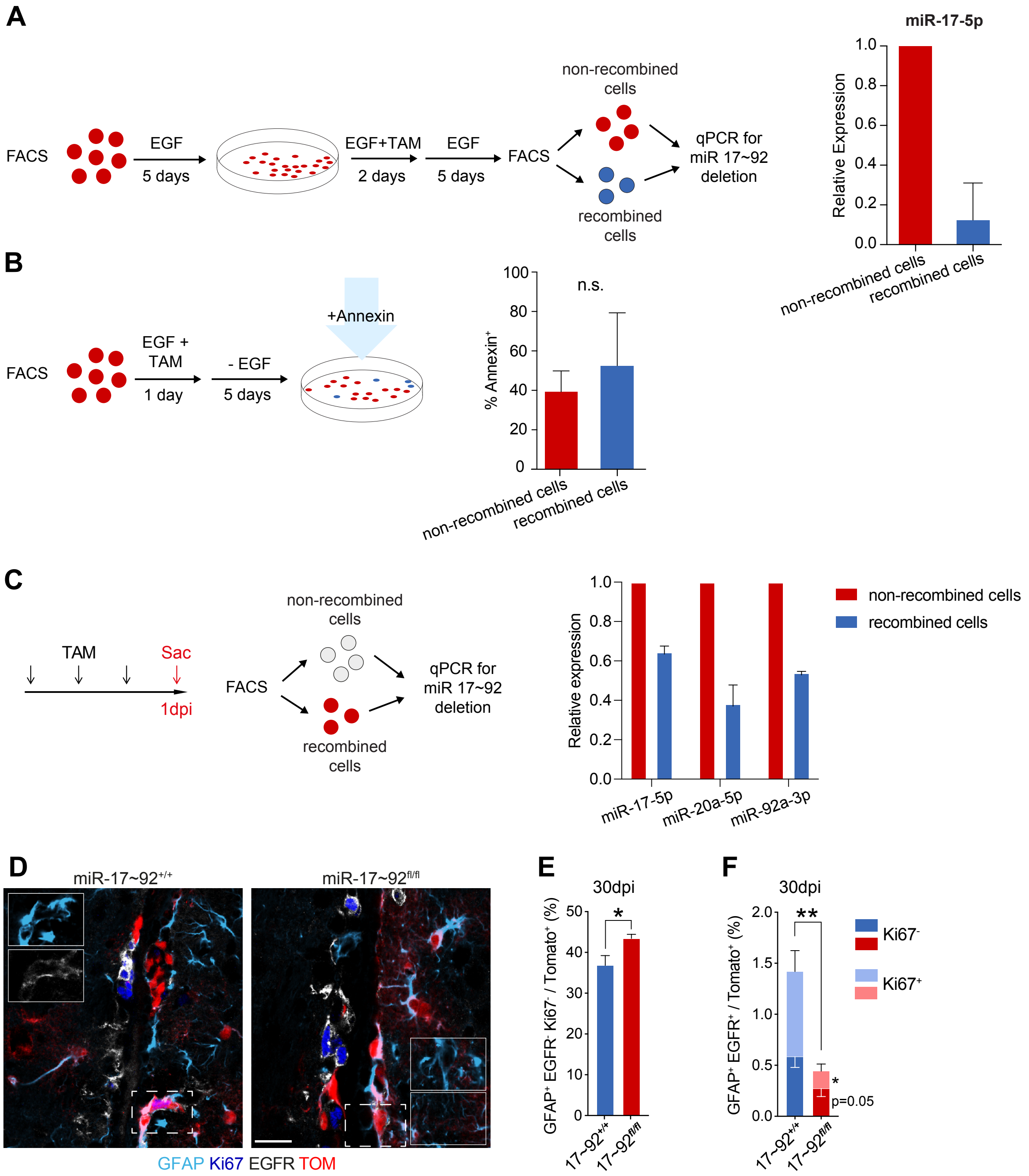

### *Supplementary Figure 3*

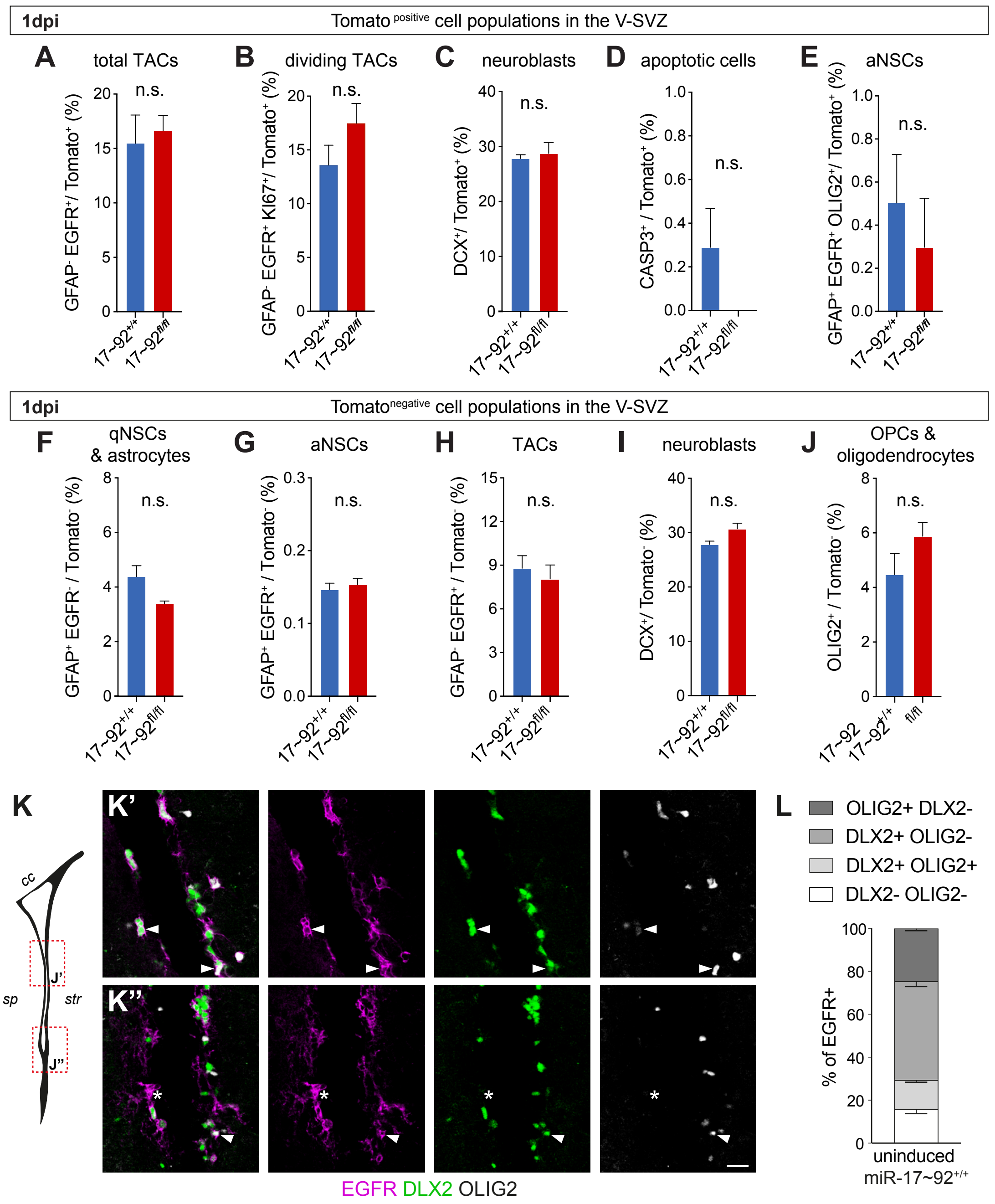

### *Supplementary Figure 4*

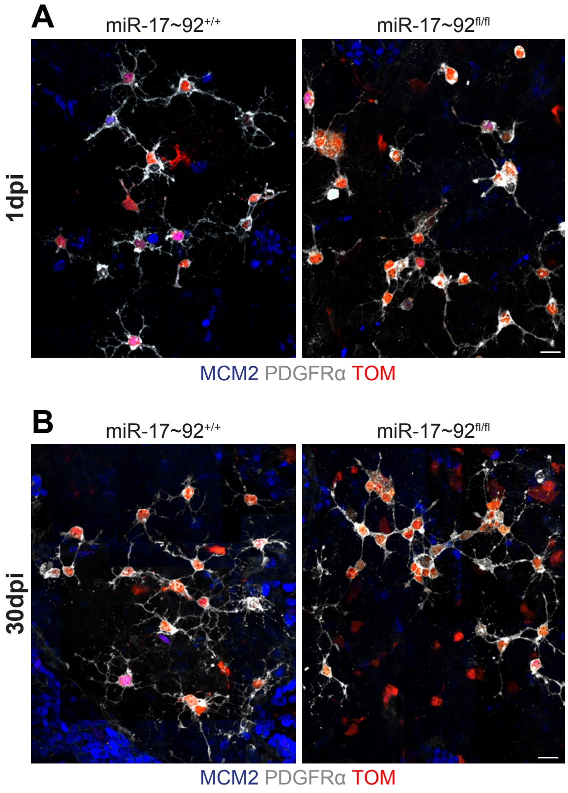
